# MetroNome — a visual data exploration platform for integrating human genotypic and phenotypic data across diseases

**DOI:** 10.1101/769646

**Authors:** Christian Stolte, Kevin Shi, Nina Lapchyk, Nathaniel Novod, Avinash Abhyankar, Lyle W. Ostrow, Hemali Phatnani, Toby Bloom

## Abstract

MetroNome is a web-based visual data exploration platform which integrates de-identified genomic, transcriptomic, and phenotypic data sets. Users can define and compare cohorts constructed from multimodal data and share the data and analyses with outside tools. MetroNome’s interactive visualization and analysis tools allow researchers to quickly form and explore novel hypotheses. The deidentified data is linked back to the source biosample inventories in multiple biobanks, enabling researchers to further investigate new ideas using the most relevant samples.

## Introduction

Biomedical research is producing a wealth of genomic data, some of it public [1]; though much is restricted to various consortia or project team members. The restrictions are often necessary to comply with consents and regulatory policies. Analyses of complex multimodal patient-derived data—such as genome sequencing, clinical and pathological measures, environmental factors, and imaging—enables questions to be explored that would otherwise not be possible. For example, how can the same chromatin remodeling genes be associated with autism, schizophrenia, bipolar disorder, congenital heart defects, and digestive tract issues? To achieve adequate statistical power for genomic research discoveries, different types of data — from different studies and diseases — must be integrated while assuring regulatory compliance with patient confidentiality and data use policies [2, 3, 4, 6]. The associated tools should be openly available and usable by a wide audience with different levels of expertise in genomics and biostatistics, while still ensuring responsible use of the data. Existing applications make it possible to integrate GWAS results with other data to prioritize variants by phenotype [7] or browse available individual-level genotype and sequence data associated with phenotypic features [8]. However, these tools generally lack the capability to then generate lists of samples and subjects matching genomic and phenotypic criteria of interest, enabling access to the underlying de-identified data and samples across multiple biorepositories.

MetroNome comprises a web-based suite of interactive data visualization tools, enabling users to combine their own data with other relevant public datasets and explore the results via linked graphs and diagrams. The system leverages human visuospatial cognitive abilities to reveal patterns and find connections. Researchers can segregate results along multiple dimensions, such as tissue source, quality control measures, whether specific variants are present, or any combination of ranges and categories in phenotypic and demographic dimensions. This enables easy comparisons between user-defined cohorts via the visualization modules. Normalized analyses, such as gene expression z-scores, are calculated in real time, based on the user’s search parameters, to allow side-by-side visual comparisons which highlight critical differences between groups, e.g., cases and controls.

## Background

MetroNome derives from two thus far distinct paths in data exploration. Genomic visualization tools such as cBioPortal [14] provide domain-specific visualizations such as variant diagrams and the oncoprint visualization. An earlier and orthogonal stream of work was the development of commercial “OnLine Analytic Processing” (OLAP) tools, which date to the 1970’s but gained traction in the 1990’s with tools such as Cognos (now IBM) [www.ibm.com/Cognos/Analytics], and Business Objects (now SAP) [www.businessobjects.com/]. These tools enabled multi-dimensional analysis of data, and dynamic filtering of commercial data along individual dimensions – although they were primarily tabular, not graphic. The second generation of OLAP, led by Tableau [www.tableau.com] introduced graphics and dashboards. Dashboards provide multiple visualizations on the same screen and the ability to filter on any one frame and propagate that filter to all other frames on the dashboard. This dynamic queries technique initially arose as an alternative to SQL for querying databases [5]. MetroNome unites these two directions: scientific and statistical visualizations, combined with dynamic filtering and filter propagation.

## Exploration of synthetic cohorts via multi-modal interactive data visualization

MetroNome presents genomic and transcriptomic data in the context of phenotypic attributes, relying on customized linked visualizations to enable exploration.

### Creating synthetic cohorts

We allow the user to combine and display data from multiple sources, based on phenotypic or genotypic traits and user access rights to those datasets. We provide access to publicly available reference datasets, such as 1000 genomes [12] and TCGA tumor data [13] for use as comparators. To perform an analysis, the user starts by creating a query that selects data from one or multiple sources. MetroNome’s query page presents a series of dynamically linked drop-down menus that can be combined into rules for selecting subjects. Rules for subject and sample criteria can include filters for any information available in the datasets that the user has selected for inclusion. Genomic rules can include the presence of variants in a list of genes or in a genomic region. Variants can be filtered by their predicted protein-coding impact as calculated by SnpEff [9], or their association with disease as recorded in ClinVar [10]. The search produces lists of subjects, samples, and variants that match the selected rules.

### Multi-modal visualization and refinement of cohorts

The query results display multiple types of data simultaneously in linked panels, such as whole genome or exome variants, RNA expression, copy number variations, structural variants, and phenotypic data, to facilitate visual exploration of associations among different data types. The visualization controls enable users to refine queries in an intuitive and dynamic fashion while exploring relationships in the data. Researchers can alter results along multiple search dimensions— for subject, sample, and genomic criteria — without rerunning their query — by refining the visualizations to samples with specific variants, or combinations of ranges and categories in phenotypic and demographic dimensions. As the user changes the extent of any one category, that change is projected to all other displayed data.

Scientists can use their intuition to generate hypotheses, then quickly look for initial confirmation, and readily refine the direction of their search to pursue a suspected causal effect. The resulting synthetic cohorts resolve to a set of individual subjects and samples, and the contents of that set can be extracted for further analysis. A frequent current use case comes from our collaboration with the Target ALS Multicenter Human Postmortem Tissue Core [11]: ALS researchers can identify decedent biosamples of specific interest to their research using MetroNome’s data exploration capabilities, and then work with Target ALS Core directors to rapidly obtain the specific blinded sample sets culled from multiple academic centers necessary for rigorous follow-up experiments.

### Comparison of cohorts

We provide the ability to view two cohorts side-by-side, to allow comparisons of traits that might influence results and warrant further study. Visually comparing datasets is one way to determine whether given cohorts are of interest, and whether specific dynamic filters better isolate features of interest. Cohort-normalized values, such as gene expression z-scores, are recalculated on the fly, based on the user’s search parameters, to accurately represent critical differences between groups, e.g., cases and controls.

### Example use case

To illustrate the utility of MetroNome, we can use examples from ALS research: Figure 1 is a comparison between patients with (left) and without (right) C9ORF72 repeat expansions, showing gene expression patterns for the gene FIG4. The neuro axis diagrams present clear differences between the two groups in the primary motor cortex and, for the cohort with repeat expansions (left), the motor cortex vs. occipital cortex, an uninvolved region. In the relationship diagrams, this cohort is also marked by shorter duration of the disease.

**Figure 1:**
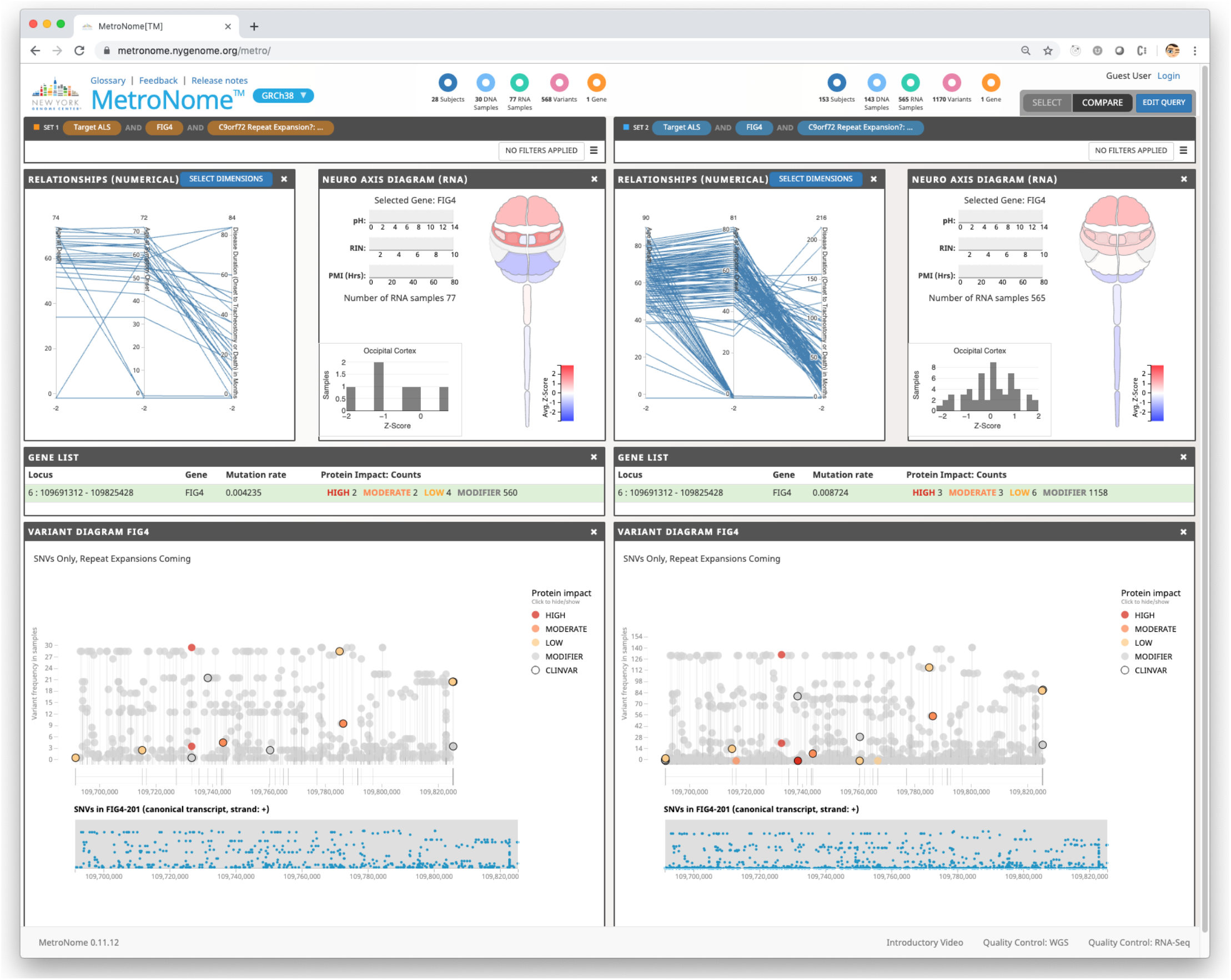
Comparison of cohorts with C9ORF72 repeat expansions (left) and without (right), showing gene expression patterns for the gene FIG4 in an anotomogram (top), and variants (bottom).

Figure 2 shows a query for samples with variants in the gene PFN1, requested by a researcher who had seen white matter abnormalities in mouse spinal cords. The neuro axis diagram clearly shows higher PFN1 expression in the spinal cord, and particularly in the thoracic spinal cord, which is interesting because the thoracic cord has a higher proportion of white matter compared to cervical and lumbar. In addition, the RNA heatmap indicates that there are a couple of specific samples with very high cortical PFN1 expression. These specific decedent tissues and slides can be selected for further benchtop experiments, and correlation with phenotypic information.

**Figure 2:**
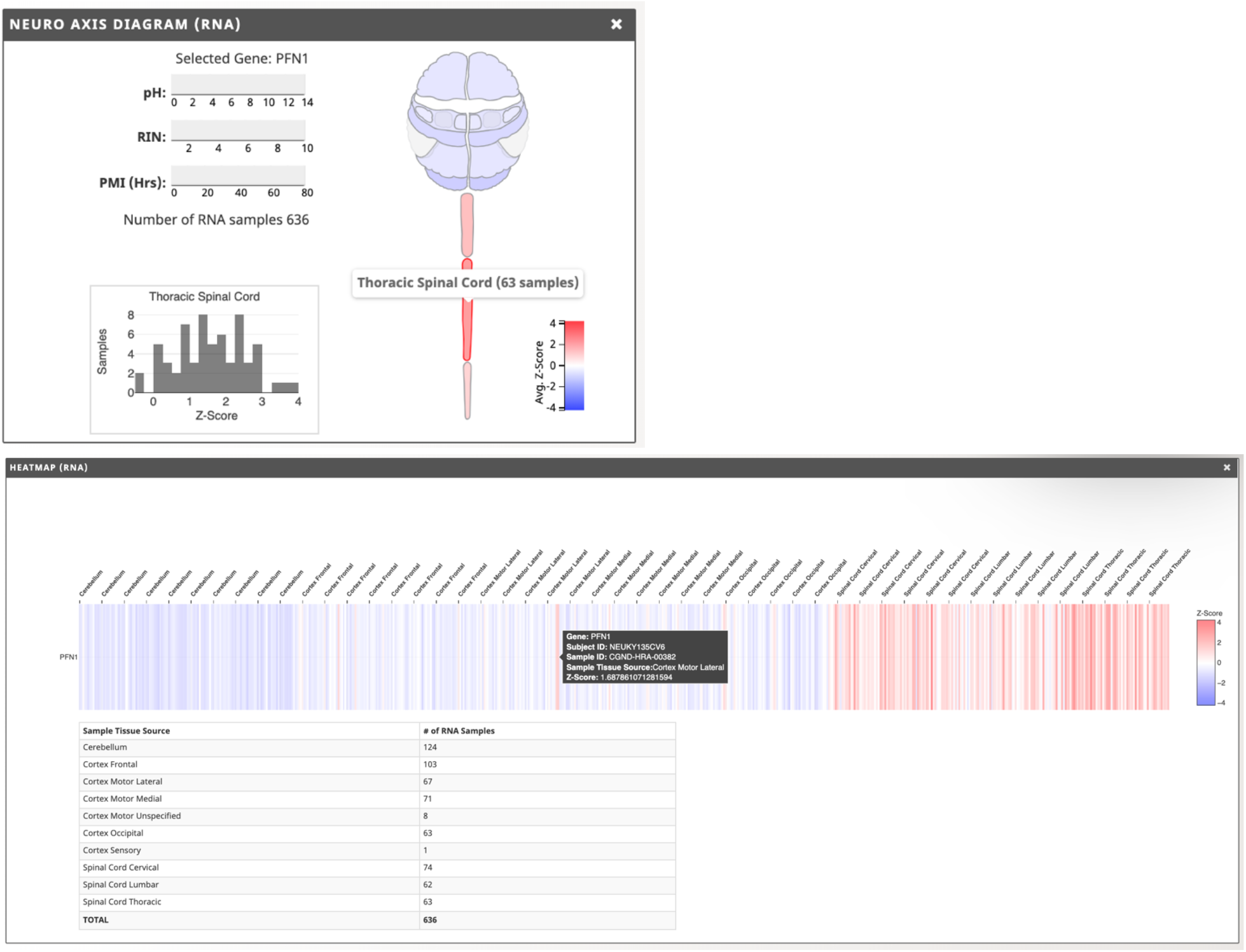
a query for samples with variants in the gene PFN1. The neuro axis diagram (top) clearly shows higher PFN1 expression in the spinal cord, and particularly in the thoracic spinal cord. The RNA heatmap (bottom), besides showing generally higher expression in the spinal cord samples, reveals a couple of samples with very high cortical levels.

These types of searches can be performed instantly and without the need for prior bioinformatics training. Lists of subjects, samples, and variants can be downloaded for further analysis and used to identify and request tissues/biofluids/slides meeting specific criteria (such as gene expression patterns, QC measures, or specific variants) from participating biorepositories for further benchtop experimentation.

## Infrastructure to support cross-study dynamic visualization

In addition to the visualization tools themselves, a great deal of data infrastructure is needed to realize the outlined goals. These infrastructure tools center on integrating data with available standards and with other data across multiple studies. We outline here three areas we have found necessary to address:

- **Data harmonization:**We harmonize data where possible, which is currently a manual process. Fields with different names in different datasets must be changed to the same standard names in each. The values of those fields must be converted to the same vocabulary, the same numeric ranges, and the same units. Values that are in reference to a specified or assumed range, such as values from some laboratory tests that have varying ranges by instrument used, must be recorded along with the reference range. Missing values must be handled, whenever possible, without discarding the entire record.
- **Use of metadata and provenance**: the data needed for analyzing results for a single study are often insufficient for integrating that data with information from other sources. Where the metadata and provenance data exist, we use such information to more accurately present combined data. We flag uncertainties to minimize misleading results.
- **Reference data**: to enable as much data integration as possible, we maintain significant types of reference data, including target sets for standard exome kits. ClinVar [clinvar.org] annotations are used to filter for pathogenic variants. Ensembl transcript-level and protein domain annotations provide information on protein-coding impact and high-impact variants, when disease significance is still unknown. 1,000 Genomes and TCGA somatic data are maintained, primarily as sources of additional data for sparse datasets. These reference data are used for comparisons, for interpretation of metadata, and for annotation of the synthesized cohorts generated in MetroNome. Note that full data integration to enable further analysis is a much more extensive problem that we have yet to address. Our work here is focused on enabling integrated and comparative visualizations.

## Privacy and security

Privacy and security are major concerns when we are supporting limited-access datasets. Authorization to access a particular dataset is determined by the owner of that dataset. The NYGC Data Privacy Committee must review the owner’s approval before access is granted within MetroNome. While this is a somewhat burdensome process, we feel it necessary to ensure that we can host private data without risk of unintentional disclosure. If a user is approved for access, they can grant members of their lab further access without review. This last case happens most often when a user is part of a consortium, and the official owner of the data is their institution. The one exception to that rule is that we allow users to upload any dataset to which they already have access, limited to personal access only, and combine it with other MetroNome datasets they can access.

The MetroNome backend adds a clause to every database query to restrict the query to the set of samples to which the user has access. Currently, users can access public data without logging in. If they do so, the backend defaults to the “public” user, with associated access rights, and limits all queries to public data only. Thus, no query or request can bypass the front end and avoid the privacy checks.

The MetroNome front-end runs in an isolated subnet, accessible from outside the firewall. The database and middle tier run behind the firewall, and all data thus resides internally. There is a single connection through that firewall, restricted to a single machine address. Authentication is performed for each connection established.

## Technical implementation

### Architecture and technologies used

MetroNome is built on a column-oriented database, Vertica, that holds all data (variants, gene expression, phenotypic, demographic, and sample-related data). An application server, written in Java, processes requests to the database and prepares results for display in the UI. The frontend is written in JavaScript, using the React framework and D3 visualization library, and is hosted in a TomCat web server.

### System requirements

The system is designed to operate on Linux virtual machine nodes running CentOS 7 and can be scaled to meet demands. The current database, Vertica, requires a cluster of dedicated nodes with matching specifications for best operation; that is, each node of the cluster should be similar in CPU, clock speed, number of cores, memory, and operating system version.

### Source code and availability

The source code is currently being converted to open source and will be made available on GitHub: https://github.com/nygenome/metronome.

The MetroNome installation hosted at the New York Genome Center is publicly available at https://metronome.nygenome.org

### Current use: Target ALS

The Target ALS Resource Cores were conceived to accelerate ALS therapy development by providing the necessary highly curated biosamples and data resources broadly and rapidly to the entire ALS research community. Given the numerous failures in translating laboratory results into clinically effective therapies, a crucial aim was to address the substantial unmet need for high quality patient-derived biosamples – such as brain, spinal cord, and muscle tissue samples from patients who died from ALS and controls, and biofluids and stem-cell lines collected during disease progression.

We perform centralized Whole Genome Sequencing (WGS), and RNA-Seq for multiple central nervous system regions at the New York Genome Center on every autopsy performed at one of the academic centers comprising the federated Target ALS Postmortem Tissue Core. After passing QC, the clinically annotated genomic and transcriptomic data is ingested into MetroNome and remains linked to the tissue samples and de-identified metadata via Global Unique Identifiers (GUIDs). The WGS and RNA-Seq raw data files (in multiple formats) are also made immediately available without embargo or IP concerns – via an online form and established data transfer workflow.

MetroNome enables researchers with very little background in genetic analysis to access the data set in a meaningful way, with relevance to their personal research interests. When researchers are interested in a specific pathway or target, they can use MetroNome to search for cases with specific mutations or variants, explore new hypotheses by comparing different cohorts defined by anatomical regions or subject groups, and then directly request tissue samples from those cases. For virtually every research project utilizing human biosamples, MetroNome can be used to refine slide and sample sets, and direct further analysis. As examples, MetroNome can be used to

- identify relevant tissue samples or slides for further benchtop experiments;
- find variants or RNA expression changes in specific targets;
- provide “clean controls” that do not possess mutations or unknown variants;
- compare spatial expression patterns with published imaging biomarker data meant to quantify relevant pathways;
- examine whether gene expression patterns are consistent with activation of pathways modulated by potential new drug candidates;
- identify whether specific patient subgroups display gene signatures that might inform patient selection for clinical trials (Figure 3);
- segregate patients based on spatial gene expression patterns and correlate with fast/slow progressors, site of onset, or specific neuropathological metadata;
- design further collaborative analysis of the genetic raw data and samples, such as whether subject groups with distinct genetic patterns might correlate with biomarker profiles in fluids or peripheral tissues.

**Figure 3:**
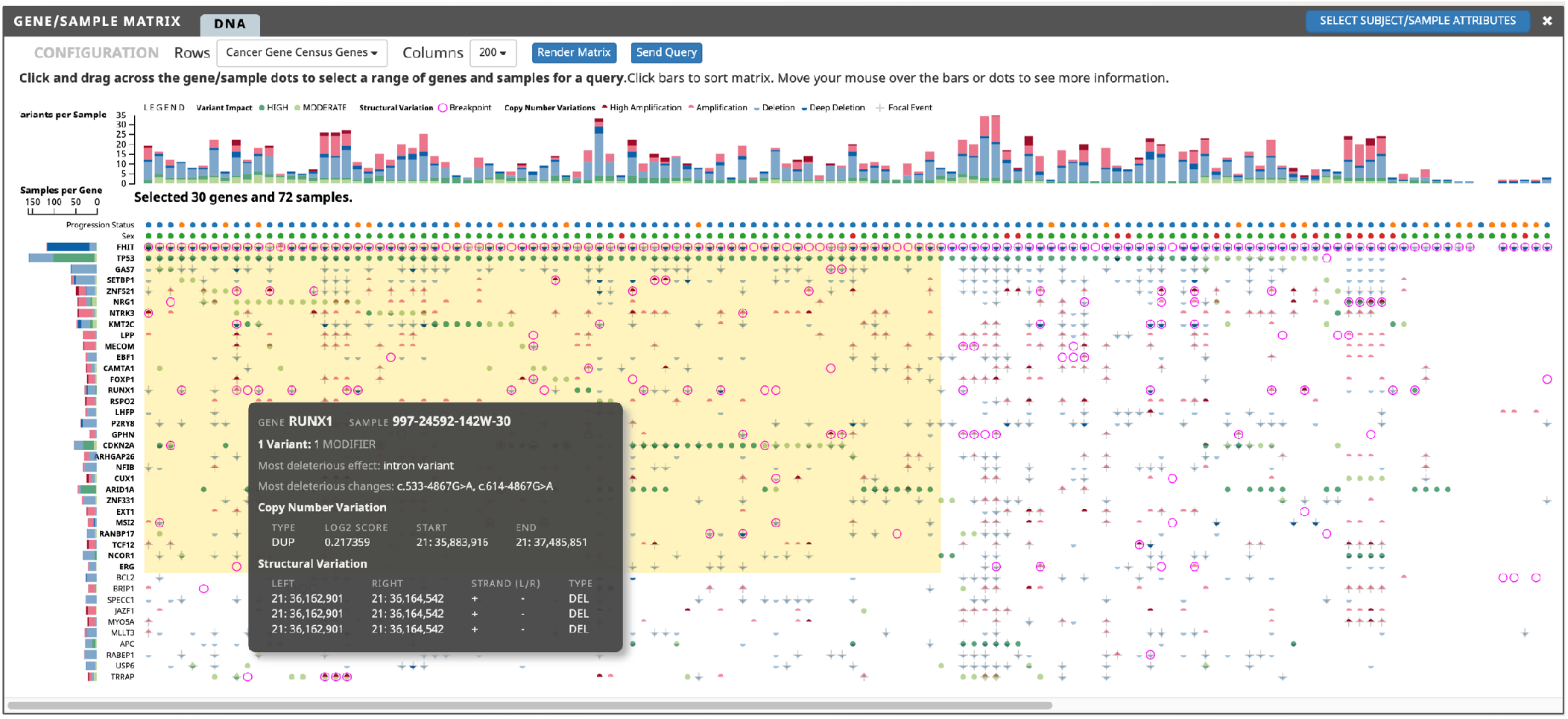
Sortable gene/sample matrix for genes from the Cancer Gene Census, shown in rows; columns represent samples. The grey callout shows details for one gene in one sample. Single-nucleotide variants are identified as high or moderate impact. Copy number changes and structural variants are identified by special glyphs and colors. Sorted by default to show genes and samples with the most impactful variants, this matrix can be used to select samples and genes (yellow area), e.g., to create a synthetic cohort prioritizing highly mutated samples.

The MetroNome visual data exploration platform has proven critical to the continued success, expansion, and evolution of the Target ALS Postmortem Core and associated efforts. It has supported over 100 different academic and industry labs, facilitating more than 135 different ALS research projects in 16 countries across 4 continents. This often includes multiple different projects in each lab. MetroNome has become part of a scientific ecosystem that includes clinics, research labs, and industry.

### General use

MetroNome is unbiased towards specific disease areas and can accommodate genomic and phenotypic data from any study. Figure 3 shows an example from an esophageal cancer study, indicating presence of single-nucleotide variants, copy number variations, and structural variations for a matrix of genes and samples.

### Future work

New releases will include some additional critical features:

- Download images from our visualizations, along with the metadata about the cohort being used and the filters applied.
- Automate the process for users to upload their own datasets, visible only to them.
- Improve reproducibility: users will be able to save queries, to rerun them in a later session, or share them with other users. The ability to save *results,* and access them later, should underlying data change in the interim, is planned. Finally, tracking the steps a user executes in a session, displaying that history and allowing a user to return to a previous state is a feature we believe to be very useful in this context.
- Add support and visualizations for new data types, such as repeat expansions and splice junctions. We expect the types of data to expand continually.
- Automate harmonization using HPO terms [15].
- Pedigree relationships, including flagging filtered de novo or recessive homozygous variants in the probands. Our structure allows subjects to be considered probands in one study and relatives in others.
- To aid evaluation of results, we want to integrate statistical analysis tools. We may link out to R, to BioConductor tools [www.bioconductor.org], or to tools such as DeepSea [16].
- APIs to allow MetroNome to exchange both data and compute with other repositories. These APIs are essential to users who wish to use the MetroNome resources with their automated analyses, rather than through our visualization interface.
- Develop user interfaces for longitudinal data

## Summary

Synthesizing cohorts by integrating data from multiple studies presents numerous challenges. Providing this functionality as part of an interactive phenotype-genotype visualization platform enables data integration as a fundamental part of the platform. Not only does this approach enhance integrated multi-modal analysis, it provides a framework that reduces the work each researcher must perform to obtain a clean cohort that meets their research needs. Using this visual data integration platform to generate and explore hypotheses is a further important contribution, with the potential to accelerate the work of scientists anywhere by eliminating the bioinformatics bottleneck during genesis of ideas. Researchers can then follow up *only the best leads* with their computational colleagues for thorough analysis.

## Acknowledgements

The authors would like to thank the Target ALS Multicenter Postmortem Tissue Core for making their sample data publicly available, and for valuable input during development of functionality in MetroNome; Kanika Arora and Minita Shah for evaluating MetroNome’s utility for cancer studies and for providing extensive feedback during development of the gene/sample matrix; Dorian Leary, Dimitrije Jevremoviç, Joseph Mulvaney, and Sylvestre Gug for their software engineering; thanks to Phaedra Agius, Michael Zody, Simon Tavaré, and Tom Maniatis for reviewing the manuscript.

